# Correction of multiple splicing mutations associated with CFTR exon 18 using a single exon-specific U1 snRNA

**DOI:** 10.64898/2026.01.30.702834

**Authors:** Annabelle G Elsner Pacheco, Jenell M Betts, Shuyu Hao, Hannah Callejas, Karl Mader, Abdul McKinney, Ronald A Conlon, Mitchell L Drumm, Hua Lou

**Affiliations:** Department of Genetics and Genome Sciences, Case Western Reserve University, School of Medicine; Department of Biochemistry, Case Western Reserve University, School of Medicine; Center for RNA Science and Therapeutics, Case Western Reserve University, School of Medicine

**Keywords:** CFTR, splicing, mutations, U1 snRNA

## Abstract

Splice site mutations represent a major class of pathogenic mutations in many diseases, as these changes disrupt normal splicing leading to gene expression changes. Cystic fibrosis (CF) results from mutations to the cystic fibrosis transmembrane conductance regulator (CFTR) gene that encodes an essential ion channel. Approximately 13% of the over 2,100 known CFTR mutations disrupt 3’ or 5’ splice sites and are predicted to cause splicing defects. Because each splicing mutation is rare, developing individualized therapies to treat each one is financially challenging. Exon specific U1 snRNA (ExSpeU1) targets the non-conserved intronic region downstream the 5’ splice site (ss) to rescue exon skipping. Because this approach is exon-rather than mutation-specific, a single agent can potentially rescue multiple mutations. In this study, we have developed a platform to systematically classify all patient variants associated with an exon that are predicted to affect splicing and then determine their rescue potential using ExSpeU1. Here we report the results of these studies. Our minigene reporter study shows that 7 of 10 exon 18 variants resulted in exon skipping. Four mutations at the 3’ and 5’ ss were rescued at least partially using a single ExSpeU1. Using a luciferase reporter, we observe that the splicing rescue is reflected at the protein level. Lastly, we demonstrate exon-targeting ExSpeU1s can also rescue 3’ and 5’ ss mutations. Overall, this study exemplifies the power of our platform to screen and rescue multiple patient-derived splicing mutations using a single agent.

## INTRODUCTION

In eukaryotes, pre-mRNA molecules undergo several processing steps in the nucleus, including capping, splicing and polyadenylation, to generate mRNA transcripts that are transported to the cytoplasm for protein synthesis. Pre-mRNA splicing removes non-coding, intervening sequences, or introns, joining coding sequences, or exons, to generate mature mRNA. This critical step in gene expression is conducted by the spliceosome, a large ribonucleoprotein (RNP) complex involving more than 100 proteins; there are five small nuclear RNPs (snRNPs): U1, U2, U4, U5, and U6 (Matera & Wang, 2014). The first step of splicing and spliceosome assembly is the binding of U1 snRNA, the RNA molecule in U1 snRNP, to the 5’ splice site (5’ ss) of a transcript through a base pairing interaction. This interaction initiates a step-by-step formation of different spliceosome complexes leading to assembly of the catalytically competent complex, complex C, which carries out the splicing reactions (Wilkinson et al., 2021; Kastner et al., 2019).

Sequence variations at the splice sites can disrupt their interactions with splicing factors such as U1 snRNA and cause aberrant splicing (Balestra, et al., 2014; Balestra, Maestri, et al., 2019). In most cases, this leads to skipping of the exon associated with the mutated 5’ ss through a process termed exon definition (Buratti et al., 2007; Leader et al., 2021; Berget, 1995). It is predicted that 15% of pathogenic variants are splicing mutations (Krawczak et al., 1992). However, this estimate only accounts for variants at the 3’ and 5’ splice sites. The actual frequency is likely much higher when including nucleotide changes that affect exonic sequences, secondary structures, and splicing enhancers or silencers (Soemedi et al., 2017; Wang & Cooper, 2007).

Splicing mutations occurring at the 5’ ss disrupt the 5’ ss-U1 snRNA base pairing interaction, often leading to skipping of the exon associated with the 5’ ss. This splicing defect can be rescued using engineered U1 snRNA that restore base pairing. Over the last 15 years, two generations of modified U1s have emerged, both of which involve altering the 5’ end of the U1 snRNA to restore base pairing.

The first generation of modified U1 snRNAs, termed suppressor U1 (SupU1), contains compensatory mutations at their 5’-ends that restore base pairing with the mutated 5’ ss. This approach targets specific pathogenic mutations, and has shown success in several minigene reporters modeling reported patient variants from several diseases including Coagulation Factor Deficiencies (Pinotti et al., 2008), Neurofibromatosis Type 1 (Baralle et al., 2003), and Tyrosinaemia Type 1 (Scalet et al., 2018). This approach has been used *in vivo* in mouse models for Factor VII (FVII) Deficiency, Tyrosinemia Type 1, and Aromatic L-Amino Acid Decarboxylase (AADC) Deficiency. However, these studies showed toxicity attributed to off-target effects (Balestra et al., 2014, 2020; Gonçalves et al., 2023; Lee et al., 2016).

The second generation of modified U1 snRNAs targets the non-conserved intronic regions downstream of the affected exon to restore splicing and is termed exon-specific U1 (ExSpeU1) (Gonçalves et al., 2023; Fernandez Alanis et al., 2012). ExSpeU1 has shown great potential to rescue splicing for numerous diseases in both cell and animal models including spinal muscular atrophy (SMA) (Dal Mas et al., 2015), Hemophila A and B (Tajnik et al., 2016), Familial Dysautonomia (Donadon et al., 2018; Romano et al., 2022), and Cystic Fibrosis (CF) (Donegà et al., 2020; Dujardin et al., 2011).

Compared to the first-generation approach, this approach has two major advantages. First, it has exhibited significantly reduced off-target effects. RNA-seq data from transgene SMA mouse models that also expressed ExSpeU1 showed only 12 genes had changes in global gene expression, indicating that there were very few measurable off-target effects (Rogalska et al., 2016). Second, because the ExSpeU1 targets downstream of the 5’ ss, it has the capability to increase exon definition and correct multiple mutations of the 5’ ss and mutations beyond the 5’ ss, which includes the 3’ ss and exonic regulatory elements (Fernandez Alanis et al., 2012). Essentially, the use of a single ExSpeU1 for a specific exon can potentially correct multiple mutations associated with the same exon.

Use of the ExSpeU1 approach in CF and Hemophilia A cell models provided proof-of-principle results. In the CF studies, ExSpeU1 was used to rescue splicing of a total number of 14 mutations associated with 6 CFTR exons: 5, 10, 12, 13, 16, and 18 (Donegà et al., 2020; Fernandez Alanis et al., 2012). In the Hemophilia A study, ExSpeU1 was used to rescue splicing of 9 mutations of *F8* exon 19 (exonic and 5’ ss variants) (Lombardi et al., 2021). For diseases in which thousands of mutations on the single gene have been identified such as CF, each individual exon is often associated with many splicing mutations. For example, using the CFTR mutation databases, we estimated that exon 18 is associated with at least 10 variants expected to affect splicing, only one of which was tested in the previous study (Donegà et al., 2020). To use ExSpeU1 as a therapeutic agent, it is crucial to test all potential splicing mutations associated with each exon to evaluate which exact patient variants can be corrected using this approach. Note that in this study, we refer to variants as nucleotide changes found in patients, and mutations as those confirmed to cause exon skipping and are thus pathogenic.

Our laboratory has developed a systematic approach to classify CF patient variants and correct those that cause exon skipping using a single ExSpeU1 for each exon. CF is a common genetic disorder caused by loss-of-function mutations to the CFTR gene, which encodes an important chloride channel. Approximately 13% of the over 2,100 known CFTR mutations disrupt 3′ or 5′ splice sites (Sasaki & Guo, 2018). Because each splicing mutation is rare, affecting only a few patients, developing individualized therapies is financially and logistically challenging. This study uses CFTR exon 18 as an example to develop a platform that systematically evaluates patient variants for splicing phenotypes and their potential to be rescued using a single ExSpeU1.

In this study, we tested 10 variants associated with CFTR exon 18 that are expected to affect splicing using mini-gene reporters. Of these 10 variants, nine are intronic, within the 3’ and 5’ ss, and one is located within the exon. Our minigene splicing assay shows that 7 of these variants cause exon 18 skipping. Using an ExSpeU1 approach, 5 mutations at the 5’ and 3’ ss were rescued. To test if ExSpeU1 can rescue splicing when placed on exon 18 instead of intron 18, we designed ExSpeU1s that target the exonic sequences at three different locations. Excitingly, these exon-binding ExSpeU1s were able to rescue exon skipping. To extend the results of our minigene splicing reporter system, we tested two mutations using a luciferase splicing reporter. We found that ExSpeU1 worked similarly in that context where luciferase activity was used as a functional readout.

## RESULTS

### Patient variants associated with exon 18 splice sites cause exon skipping in a minigene splicing reporter

Exon 18 is associated with ten variants found in patients in the CFTR1 database that were predicted to affect splicing because of their locations on the 5’ ss, 3’ ss, or deeper into intron 18 (**Figure 1A**). Two of the ten variants, 2909-1G>A and 2988+1G>A, are included in the CFTR2 database, which includes only the experimentally verified variants; the rest remain unclassified. The nomenclature of the variants follows the Human Genome Variation Society nomenclature in which “+” indicates the position of the nucleotide from the 5’ splice site (donor site) and “-” indicates distance from the 3’ splice site (acceptor site) (Ogino et al., 2007).

**Figure 1.**
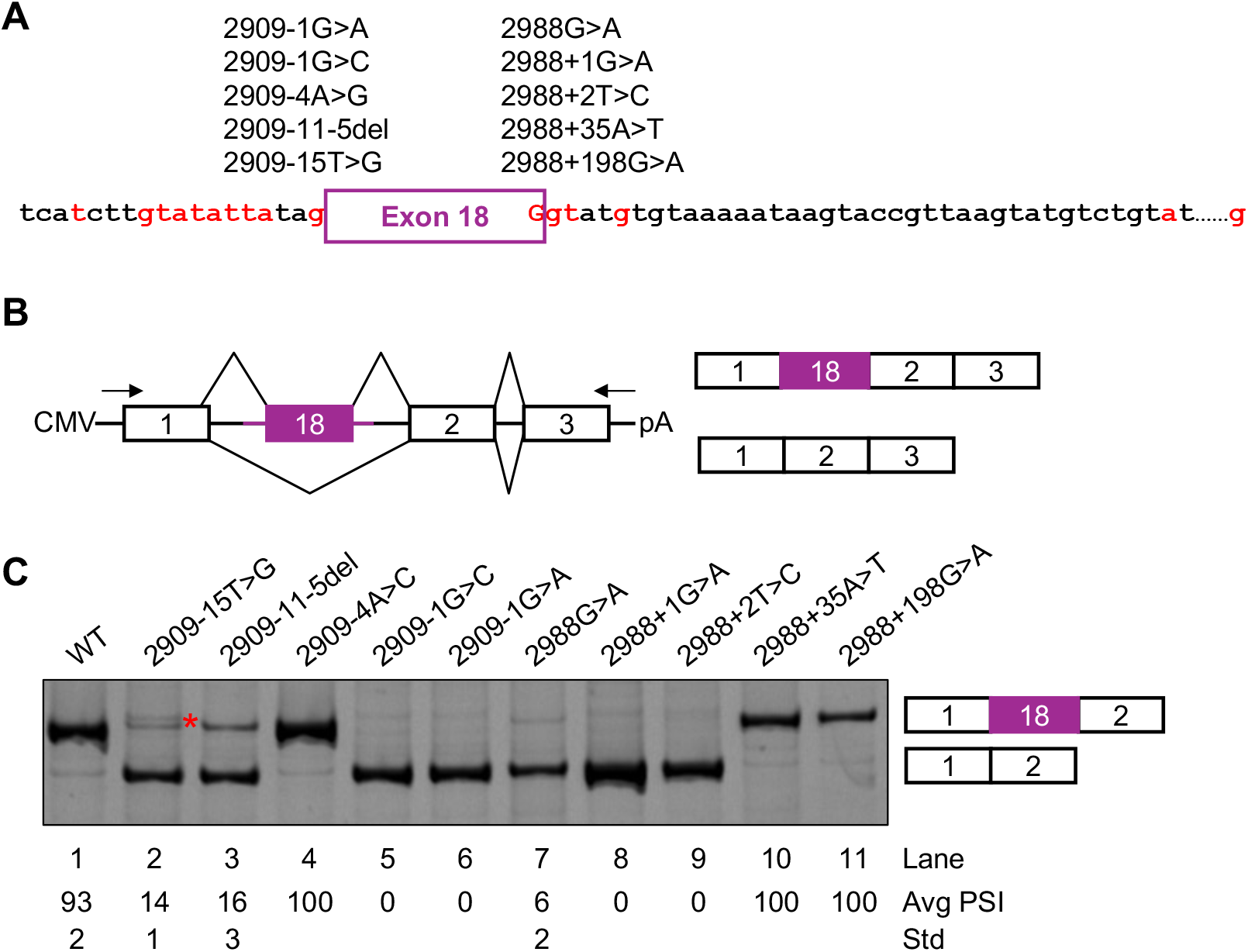
CF patient variants associated with exon 18 splice sites cause skipping of CFTR exon 18 in a minigene splicing reporter. A. Diagram of the 10 known patient variants associated with CFTR exon 18 splice sites, marked in red. There are 5 mutations at the 3’ ss (left column) and 5 mutations associated with the 5’ ss (right column). B. The minigene splicing reporter includes CFTR exon 18 and its flanking intronic regions inserted between exons 1 and 2 of the human Metallothionein 2A gene. The arrows represent primers used for RT-PCR reactions (right). C. RT-PCR analysis. Percent spliced in (PSI) and its standard deviation (Std) are shown below the gel. The red star indicates a band that results from a cryptic splicing product (see Supplemental Figure 1).

To determine the effects of each variant on the splicing pattern of exon 18, we established a splicing reporter by cloning CFTR exon 18 and its flanking intronic sequences, 305 nt of intron 17 and 335 nt of intron 18, into the human Metallothionein-2A (MT2A) gene (**Figure 1B**). Variants were introduced into this CFTR exon 18 splicing reporter through mutagenesis cloning.

The wild type and mutant exon 18 reporters were transiently transfected into 16HBE14o-cells, a human bronchial epithelial cell line. RT-PCR analysis of the isolated total RNA showed that variants 2909-15T>G, 2909-11-5del, 2909-1G>C, 2909-1G>A, 2988G>A, 2988+1G>A, and 2988+2T>C result in exon 18 skipping (**Figure 1C**, lanes 1-3, 5-9). However, the intronic variants (2988+35A>T, and 2988+198G>A) as well as 2909-4A>G, showed no skipping phenotype (**Figure 1C**, lanes 4 and 10-11). The exon 18 skipping and inclusion products were confirmed via Sanger sequencing of gel purified RT-PCR products.

An additional band slightly bigger than the inclusion band was observed for the 2909-15T>G variant indicative of cryptic splicing. This product was cloned via TA cloning and sequenced. It is 14 nucleotides longer than the correctly spliced product. As indicated in **Supplemental Figure 1**, the 2909-15T>G change created a cryptic 3’ ss and as a result, weakened the normal 3’ ss, leading to exon 18 skipping as the major splicing product.

### Rescue of exon 18 inclusion for the 2988 G>A mutant splicing reporter

The 2988G>A mutation is a silent mutation that codes for the same amino acid as the wild-type sequence. This nucleotide change occurs at the last position of exon 18 and is part of the 5’ ss that base pairs with U1 snRNA during splicing. The G>A change disrupts the base paring interaction with U1 and leads to almost complete skipping of the exon (**Figure 2A**, lane 7 in Figure 1C and lane 2 in **Figure 2B**).

**Figure 2.**
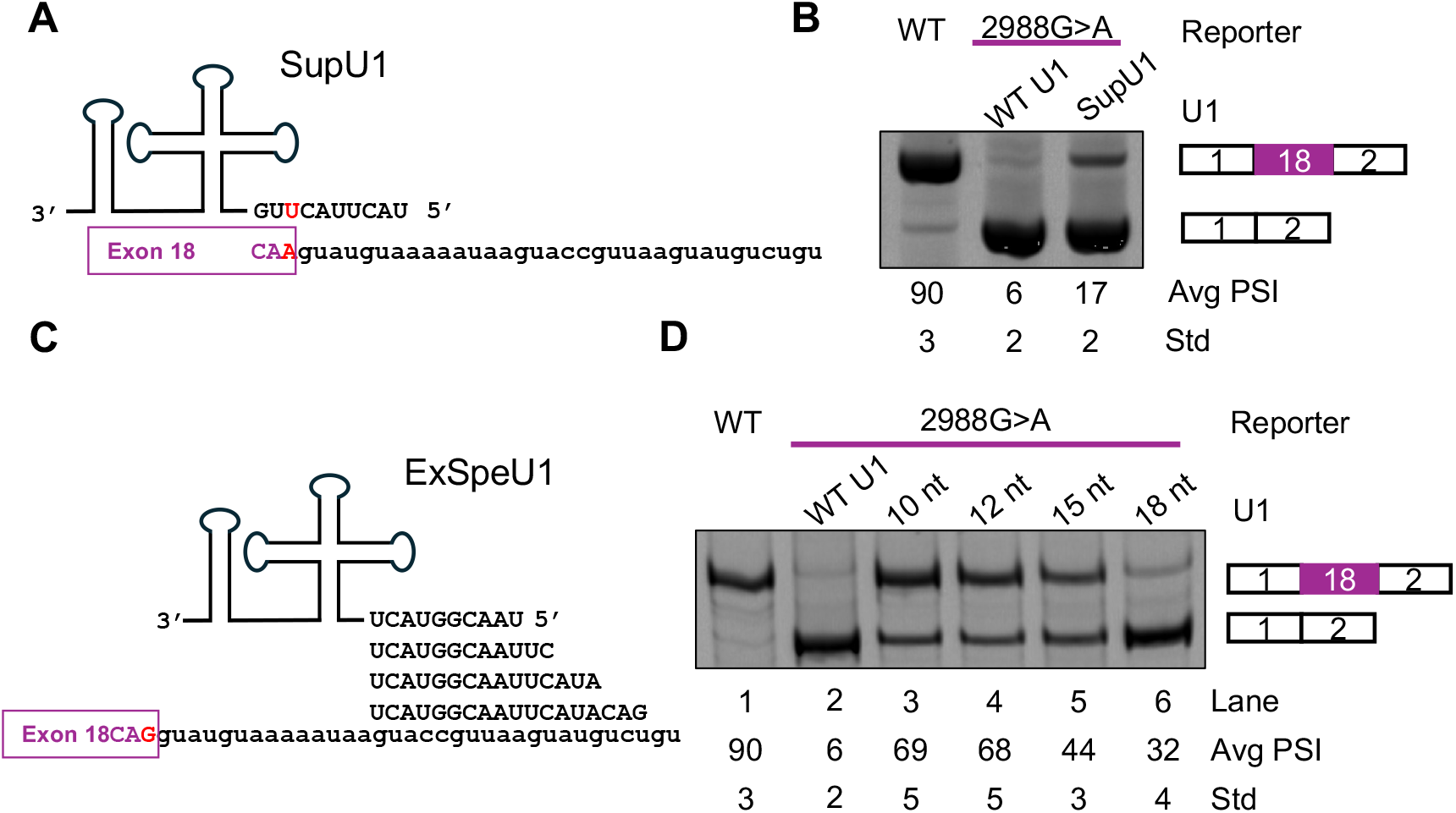
The 2988G>A mutation can be rescued using the suppressor U1 and ExSpeU1 approach. A. Diagram of binding location of Suppressor U1 (SupU1) showing compensatory mutation in red that restores base pairing to the mutated 2988G>A reporter. B. RT-PCR analysis. PSI and Std are shown below the gel. C. Diagram of the binding locations of exon-specific U1 snRNA (ExSpeU1) with varying complementary lengths. D. RT-PCR analysis of exon 18 inclusion of wild type or 2988G>A mutant reporter co-transfected with WT U1 and ExSpeU1 of complementary lengths 10, 12, 15, and 18 nt. PSI and Std of are shown below the gel.

We wished to rescue exon skipping caused by this mutation using both the SupU1 that carries the compensatory mutation, and an ExSpeU1 approach. To create the suppressor U1 a single nucleotide was changed from the wildtype U1 sequence to restore base pairing. The ExSpeU1 base pairs with intronic sequences near the 5’ ss of exon 18. As shown in **Figure 2B**, when co-transfected with the 2988G>A reporter, the SupU1 restored exon inclusion to 17%.

To evaluate modifications that would affect specificity of the ExSpeU1, we replaced the first 10 nucleotides of the U1 snRNA sequence with an anti-sense sequence that base pairs with the non-conserved intron sequence of 10, 12, 15 and 18 nt in length (**Figure 2C**). The targeted intron sequence starts at nucleotide 14 in the intron. When co-transfected with the 2988G>A reporter, all the tested ExSpeU1s increased exon 18 inclusion (**Figure 2D**). Interestingly, the ExSpeU1 with the 10 and 12 nt complementary sequences increased exon 18 inclusion with greater efficiency, at 69% and 68%, than the longer ExSpeU1s, at 44% and 32%. In subsequent experiments, we used the ExSpeU1 that has 10 nt replaced sequence at the 5’-end.

### ExSpeU1 rescues multiple 5’ and 3’ ss mutations

To systematically investigate the potential of a single ExSpeU1 to rescue several mutations associated with exon 18 skipping, we tested the approach on the seven variants that caused exon skipping (**Figure 1**). Mutant splicing reporters were co-transfected with the ExSpeU1 that base pairs with 10 nt downstream of the 5’ ss (**Figure 2**). Results shown in **Figure 3** indicate that exon skipping caused by multiple mutations at either the 5’ ss or 3’ ss were rescued by ExSpeU1.

**Figure 3.**
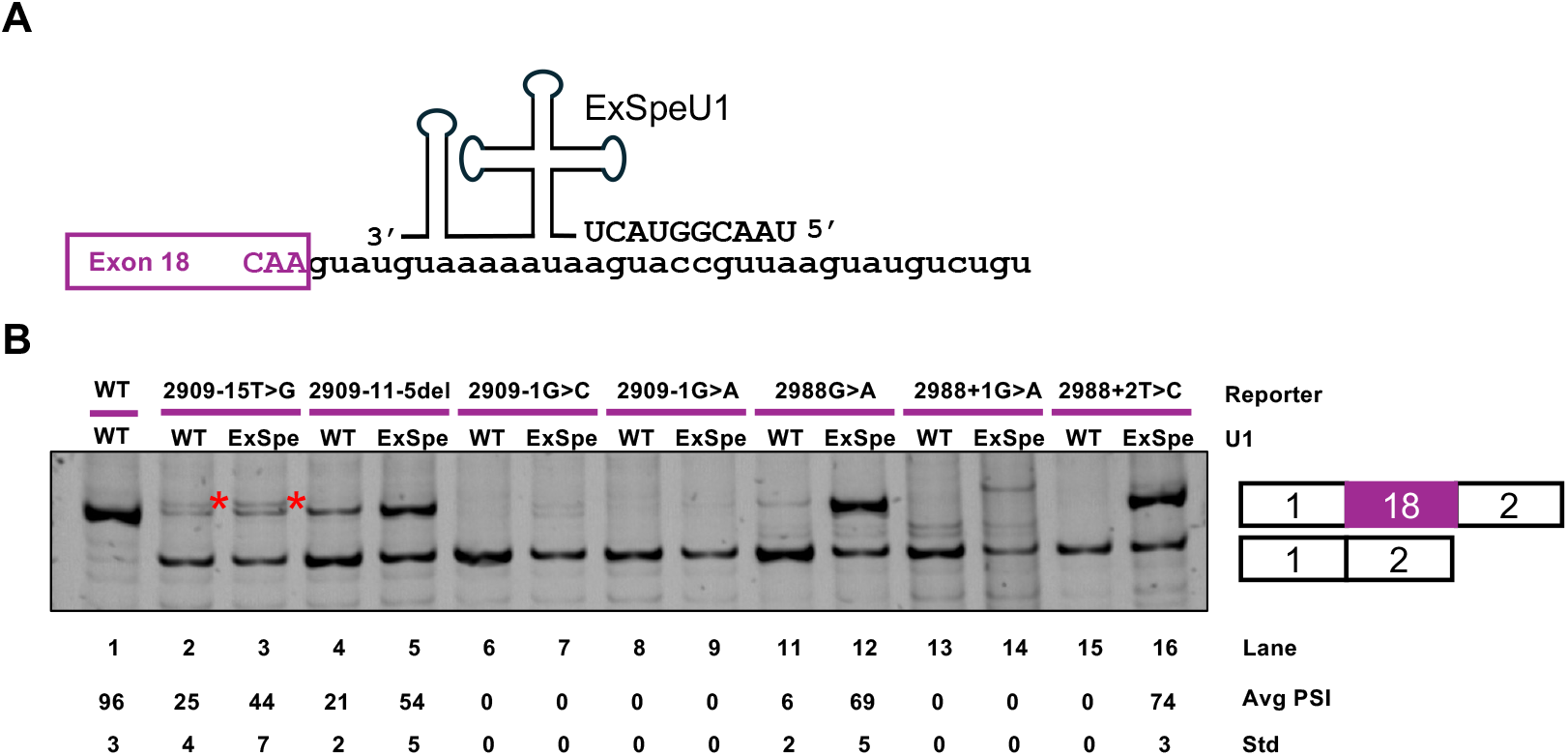
ExSpeU1 rescues multiple exon 18 5’ and 3’ ss mutations. A. A diagram depicting the binding location and composition of the complementary nucleotides of ExSpeU1. B. RT-PCR analysis to observe splicing pattern of each mutation with either WT U1 or ExSpeU1 treatment. PSI and Std are shown below the gel.

At the 5’ splice site, exon skipping of the 2988G>A mutant was rescued to 69% exon inclusion (**Figure 3**, lane 11), and the 2988+2T>C mutant to 74% (**Figure 3**, lane 15). At the 3’ ss, exon inclusion of the 2909 −15T>G mutant was increased from 25% to 44% (**Figure 3**, lanes 2 and 3), and the 2909-11-5del mutant from 21% to 54% (**Figure 3**, lanes 4 and 5).

Mutations at two positions that cause exon skipping were not rescued by ExSpeU1. These are 2988+1G>A, which occurs at the first nucleotide of the intron and mutations 2909-1G>C and 2909-1G>A, which occur at the last nucleotide of the intron (**Figure 3** lanes 7 and 9).

To optimize rescue of exon skipping due to mutations at the 3’ splice site, we conducted a dosage experiment by co-transfecting the 2909 −11-5del minigene with increasing amounts of ExSpeU1 from 0 ng to 1.5 ug. At all concentrations tested above 250 ng, we did not detect any significant changes (**Supplemental Figure 2**).

### ExSpeU1 placed in different positions within the exon can rescue 3’ and 5 splice site mutations

To explore the potential of an ExSpeU1 to strengthen exon definition weakened by mutations to the 3’ ss, ExSpeU1s were designed to bind inside of CFTR exon 18 in the splicing reporter. We tested ExSpeU1 targeting 7 (ET-7), 20 (ET-20), and 40 (ET-40) nt from the 3’ end of exon 18 (**Figure 4A**). All ExSpeU1 treatments were able to at least partially rescue exon skipping due to the 2909-15T>G and 2909-11-5del mutations (**Figure 4B**). Rescue was highest as the binding location of the ExSpeU1 approached the 5’ ss. Using the ExSpeU1 that binds only 7 nt into exon 18, exon skipping due to mutations 2909-15T>G and 2909-11-5del were rescued similarly as when rescued with the ExSpeU1 designed to bind the intronic region (**Figure 4B** lanes 2, 5, 7 and 10). As expected, none of the ExSpeU1 tested were able to rescue exon 18 skipping caused by the 2909-1G>A mutation (**Figure 4B** lanes 16-19).

**Figure 4.**
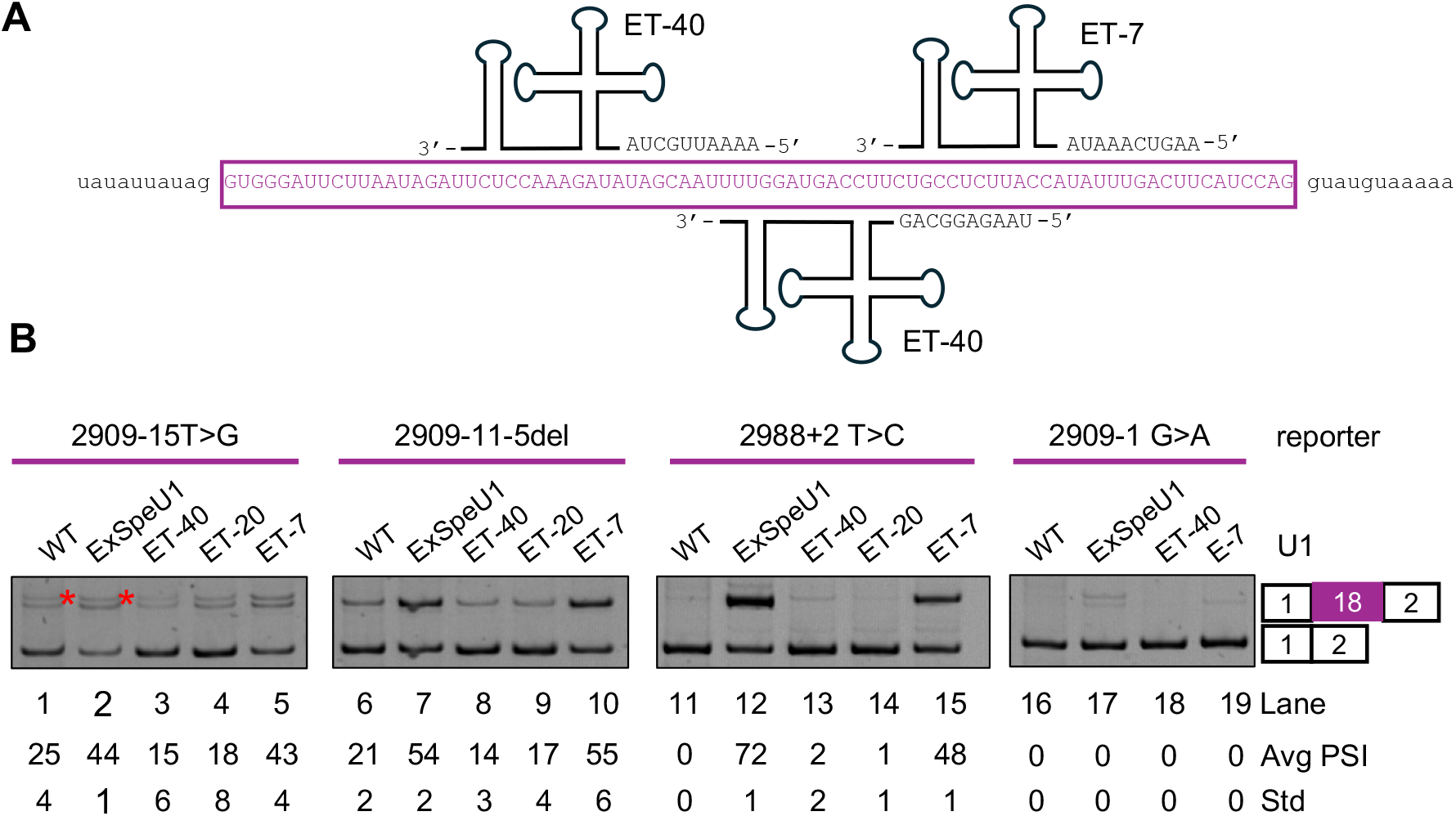
ExSpeU1 that target exonic sequences can rescue exon 18 splice site mutations. A. Diagram of binding location of ExSpeU1 that bind within exon 18 (purple). These Exon targeting ExSpeU1s bind at 7 (ET-7), 20 (ET-20), and 40 nt (ET-40) from the 5’ss. B. RT-PCR analysis. PSI is shown below the gel. A red star indicates a cryptic splicing product.

To evaluate this approach on 5’ splice site mutations, these ExSpeU1s designed to bind within exon 18 were co-transfected with the 2989+2T>C mutant reporter. While the ExSpeU1 that binds further into the exon gave very minimal rescue, the ExSpeU1 that bound only 7 nt from the 5’ ss was able to rescue exon 18 inclusion from 0 to 48%.

### ExSpeU1 strategy to correct exon 18 skipping in a fluorescent protein/luciferase splicing reporter

To determine the exon skipping rescue effects of ExSpeU1 at the protein level, we used a fluorescent protein/luciferase splicing reporter. In addition to the wild type reporter, two mutant reporters were generated, one to model the 2988G>A mutation and another the 2988+1G>A mutation. In these reporters, exon 18 and flanking intronic regions of CFTR (100 nt upstream and 75 nt downstream) were inserted between mWasabi and mScarlet/Aka luciferase (AkaLuc) (**Figure 5A**). Mutations that affect splicing and cause exon 18 skipping will result in a frameshift, reducing the mScarlet and AkaLuc activities.

**Figure 5.**
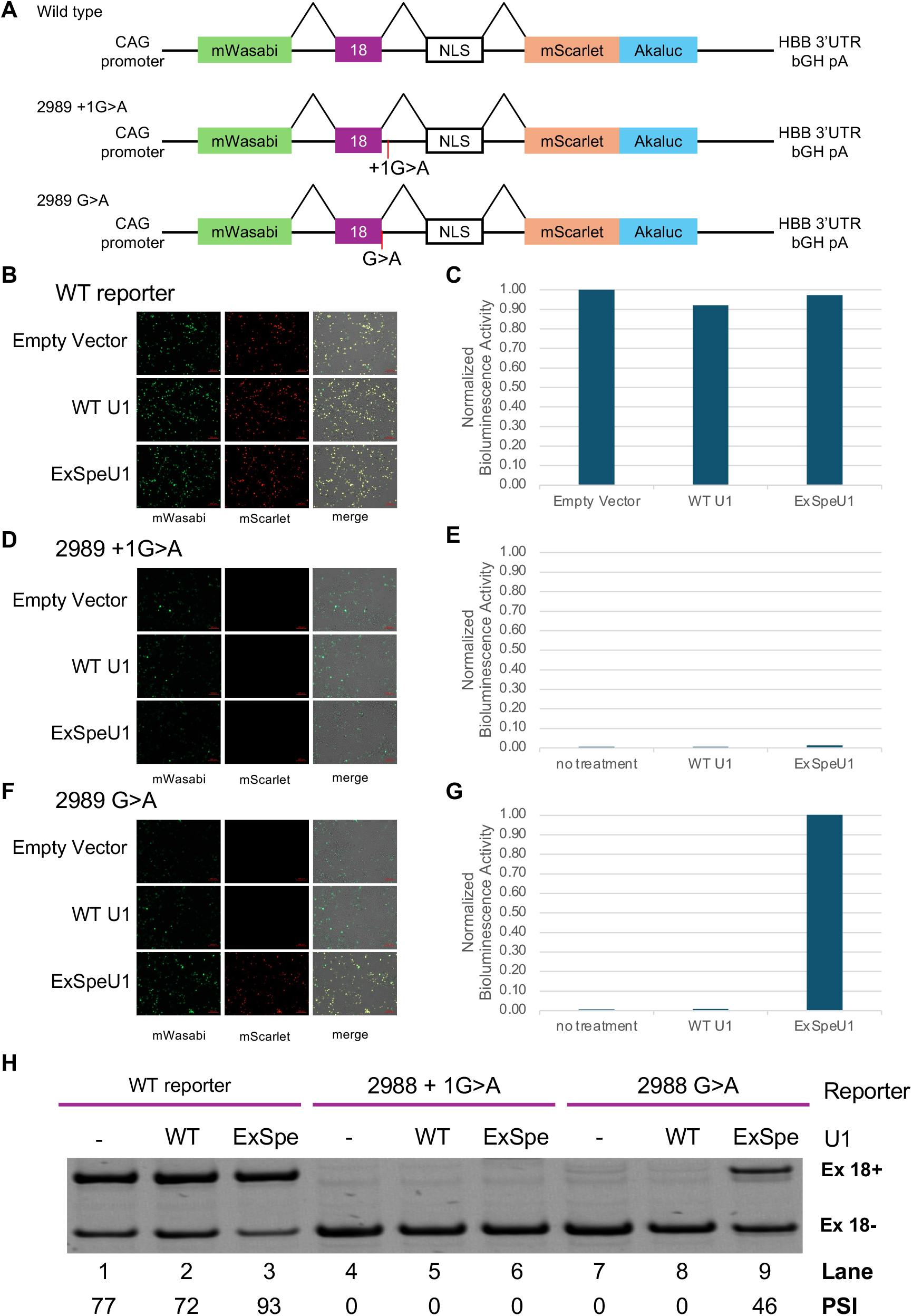
ExSpeU1 corrects exon 18 skipping in a fluorescent protein splicing reporter. Diagram of wild type and mutant fluorescent protein splicing reporter showing CFTR exon 18 and flanking intronic regions inserted between mWasabi and mScarlet/Aka luciferase (Akaluc) coding sequence. Both 2988G>A and 2988+1G>A results in frameshift mutations that reduce mScarlet and Akaluc activity. B-D. Fluorescent and bioluminescent activity of the wildtype reporter (B), 2988+1G>A (C) and 2988G>A (D) after co-transfection with empty vector, wild type (WT) U1, or ExSpeU1 in HEK293 cells. E. RT-PCR analysis. PSI is shown below the gel.

As shown in **Figure 5**, we observed a strong correlation between mScarlet/Akaluc activity and exon 18 inclusion. When transfected into HEK293 cells, both 2988G>A and 2988+1G>A reduced mScarlet and AkaLuc activities (compare top panels for the mCarlet image in **Figure 5B, 5D and 5F** and left bars in **Figure 5C, 5E and 5G**). RNA analysis showed that these mutations caused exon 18 skipping in the fluorescent/luminescent reporter (**Figure 5H**, lanes 1, 4, and 7). The same results were observed when the reporters were co-transfected with the wild-type U1 construct (compare middle panels for the mCarlet image in **Figure 5B, 5D and 5F**, middle bars in **Figure 5C, 5E and 5G** and lanes 2, 5 and 8 in **Figure 5H**). Significantly, when the 2988G>A mutant reporter was co-transfected with ExSpeU1, mScarlet and AkaLuc activities were restored (compare bottom panels for the mCarlet image in **Figure 5B, 5D and 5F** and right bars in **Figure 5C, 5E and 5G**) and exon 18 inclusion was rescued (**Figure 5H**, lanes 7 and 9). However, there was no restoration of exon 18 inclusion from the 2988+1G>A mutant reporter with the ExSpeU1 treatment (**Figure 5E**, lanes 4 and 6), consistent with the results obtained with the minigene splicing reporter (**Figure 3**)

## DISCUSSION

### Platform to study splicing mutations

Many CFTR variants that are predicted to affect splicing are rare and there are currently no available treatment options for CF patients who harbor these types of mutations. A systematic platform to classify all patient variants associated with a single exon that are predicted to affect splicing and then screen for mutations that can be rescued using a single ExSpeU1 would streamline therapeutic development.

We have developed such a platform and tested it using CFTR exon 18 in the present study. We are currently using our model to systematically test suspected splicing variants associated with other exons of CFTR and then determine which can be corrected using ExSpeU1s. Additionally, this system can be used for other monogenic diseases to screen patient variants for exon skipping phenotypes and determine their potential to be rescued. This platform, in combination with ExSpeU1s, will facilitate the identification of splicing mutations and increase treatment options for those affected by splicing mutations.

### Screening for skipping phenotype

In this study, we used CFTR exon 18 to assess the systematic approach and classify all reported patient variants associated with exon 18 that were predicted to perturb splicing because of their location near splice sites, and then determine the ability of a single ExSpeU1 specific to exon 18 to rescue exon skipping due to each mutation. Using our minigene splicing reporter, we classified all ten of the reported exon 18 variants. Seven of these variants caused exon 18 skipping. The 2909 −15 T>G mutation reduced normal splicing in multiple ways. The mutation created a cryptic 3’ splice AG at the −15 position, leading to a low-level splicing product that is 14 nt longer while decreasing usage of the wild type 3’ ss. It appears that the two competing 3’ splice sites reduced exon 18 recognition by the splicing machinery, resulting in an overall reduction of exon 18 inclusion.

The 2909 −4 A>C variant did not cause exon 18 skipping. This result is consistent with the *in silico* whole genome-wide analyses of the 3’ ss, which indicate each base is similarity represented at the −4 position, suggesting that mutations at this position of the 3’ ss would be well tolerated (Ma et al., 2015).

The variants at the +35 and +198 position into intron 18 did not cause exon 18 skipping in the minigene splicing reporters. In this region of the intron, regulatory elements, such as splicing enhancers or silencers, RNA-binding protein motifs, or secondary structures that regulate splicing may be present. Our results suggest that in the context of the minigene reporter, these two mutations are not disease-causing or pathogenic. However, the splicing reporter contains only 335 nt of intron 18. We do not rule out the possibility that in their natural context, these variants cause exon 18 skipping.

### ExSpeU1 can rescue multiple splicing mutations

After classifying the splicing phenotype of each variant, we used a single ExSpeU1 construct to rescue each variant classified as a splicing mutation. We were able to rescue exon skipping due to mutations at both the 5’ and 3’ ss. Notably, we rescued the 2988+2T>C mutant to 74% and the exonic 2988G>A mutation to 69% of exon 18 inclusion. The 2988G>A patient variant that causes CFTR exon 18 skipping was rescued in a previous report (Donegà et al., 2020). In this study, a slightly higher rescue (85%) was observed using a minigene splicing reporter that includes the CFTR sequence from exons 17 and exon 19 and two ExSpeU1s targeting intron 18 sequence starting at 10 or 13 nucleotides in the intron (Donegà et al., 2020). In addition to rescuing this mutation using an ExSpeU1 approach, we show that ExSpeU1 gives greater rescue than the SupU1, suggesting that a A-U base pair is significantly weaker than G-C base pair. Also, a base pair length of 10 nt gave the greatest rescue with this mutation, consistent with the finding that hyper-stabilized U1-5’ ss interaction impedes the switch of U1 for U6 in the subsequent step in splicing (Staley & Guthrie, 1999).

Further, the 2988 G>A variant causes the synonymous mutation, Q998Q, indicating that after splicing correction CFTR function should be restored. Importantly, if this mutation were to induce an amino acid change, this treatment in combination with the highly successful CF modulator therapy could restore CFTR levels and function by not only correcting splicing defects but also enhancing the folding, trafficking, and conductance of the ion channel.

Mutations at the +1 and −1 positions occur at the highly conserved splice site sequences and were unable to be rescued using ExSpeU1, consistent with previous reports (Balestra et al., 2020; Balestra, Giorgio, et al., 2019; Fernandez Alanis et al., 2012). In yeast, mutations to the +1 position inhibit the second step of splicing, which includes lariat cleavage and exon ligation, and can result in splicing inhibition (Horowitz, 2012; Vijayraghavan et al., 1986). Correcting the +1 and −1 positions remains a limitation of all engineered U1 approaches (Gonçalves et al., 2023).

We also used a luciferase/fluorescent protein reporter with coding luciferase sequences for the 2988G>A and 2988+1G>A splicing mutations. Using these reporters, we observed the effects of splicing rescue by ExSpeU1 at the protein level by detecting luciferase and fluorescent protein activities. As expected, fluorescence and luciferase activity was restored when the 2988G>A reporter was co-transfected with ExSpeU1.

During splicing, in addition to U1 binding to the 5’ ss, U6 snRNA also base pairs at the 5’ ss, specifically the +4 to +6 positions to coordinate the transesterification reactions of splicing. U6 does not need to base pair in all of these positions and the identity of the base-paired nucleotides does not affect splicing (Hwang & Cohen, 1996). This allows for the use of modified U6s that have been designed to carry the compensatory mutation to the +4 to +6 positions at the 5 ‘ss to rescue exon skipping, most significantly as a combined treatment with a modified U1 (Schmid et al., 2013). For the case of wild type CFTR exon 18, the identity of the +4, +5, and +6 positions are UGU meaning that U6 would base pair in all of these positions which is deemed optimal for exon definition (Hwang & Cohen, 1996). According to this model, using a modified U6 co-treatment would not increase exon definition and improve splicing for exon 18. However, if the identity of these positions were different, as they will likely be for other CFTR exons, a modified U6 co-treatment could be beneficial.

### ExSpeU1 targeting exon sequences

By designing ExSpeU1s to target exonic regions, we were able to restore exon inclusion across mutations at both the 5’ and 3’ ss. While the binding position of U1 is known to be flexible, most effort is put into designing the U1 to target downstream the 5’ss in the intron instead of placing the U1 on the exon. Here, we show that U1 can bind on the exon and still contribute to exon definition and coordinate splicing. Further, our results show that exon-binding ExSpeU1 can rescue exon skipping due to mutations at the 3’ ss with an efficiency approaching that of the ExSpeU1 targeting the intron sequence.

This finding is important in expanding the window of targetable regions of ExSpeU1 as regulatory elements can interfere with ExSpeU1 binding in the intron. Donegà and colleagues found an intronic splicing silencer, which happened to be a stem loop, downstream the 5’ss of CFTR exon 13 that prevented their ExSpeU1s that target this region from binding, limiting efficiency of correction (Donegà et al., 2020). Targeting the exon sequences rather than the intron as we have done here will widen the range in which ExSpeU1 can target to get around these regulatory regions and still rescue exon skipping.

### Future directions and conclusion

The goal of this study is to develop a platform that systematically classifies all patient variants associated with a single exon that are suspected to be splicing mutations and assesses the potential for each to be reduced using a single ExSpeU1. In this study, we used CFTR exon 18 as an example. While ExSpeU1 has shown to assist in the production of correct and functional protein, it remains important to validate these results in the expression and function of endogenous CFTR (Dal Mas et al., 2015). Additionally, the use of ExSpeU1 has shown to be safe and effective in treating mouse models for SMA (Rogalska et al., 2016). A logical next step is to validate the rescue approach on full-length endogenous CFTR and conduct functional assays on the resulting rescued protein product.

Using a single agent to rescue multiple splicing mutations in a variety of positions eases the burden for therapeutic development given that each variant is rare. Our minigene splicing reporter system offers a systematic approach to screen patient variants for mis-splicing phenotype and determines the potential for them to be rescued using a single approach that can be applied to other monogenic diseases. This is especially important as splicing mutations often cause degradation of the faulty transcripts resulting in complete lack of protein production. Given the currently used small molecules were developed to treat CF by facilitating folding, improving trafficking and increasing conductance, they do not work for patients affected with splicing mutations. An ExSpeU1 snRNA strategy provides a promising therapy for these patients.

## METHODS and MATERIALS

### Plasmid constructs

The CFTR exon 18 reporter plasmid was generated by first inserting exon 18 with 305 nt of intron 17 and 335 nt of intron 18 sequences into intron 1 of the human Metallothionein 2A (hMT2A) gene using the nested PCR approach. Templates for PCR included the RSV-HMT plasmid (Lou et al., 1994) and genomic DNA isolated from WT 16HBE14o-cells. The fragment was then inserted into the pCS2-MT plasmid (Addgene). Splicing variants were generated in the reporter using the Phusion Site-Directed Mutagenesis Kit (Thermo Scientific™ F541).

U1 snRNA expression plasmids were generated using a previously described expression unit (a gift from Dr. Joan Steitz at Yale University). Mutations were created in the U1 plasmid via site-directed mutagenesis, then the modified U1 was inserted into the pAAV-CAG-GFP plasmid (Addgene). Sequences for all reporters and U1 constructs were verified via Sanger sequencing.

The fluorescent protein splicing reporter consists of a CAG promoter (Okabe et al., 1997), and coding sequences that would generate a fusion protein of mWasabi (Ai et al., 2008), the test exon, a nuclear localization sequence, mScarlet (Bindels et al., 2017) and Akaluciferase (Iwano et al., 2018). The test exon in this case is exon 18 of human CFTR with 100 bp of the upstream intron and 75 bp of the downstream intron.

### Cell culture and transfection

16HBE14o-cells were grown in Minimum Essential Medium (MEM; Gibco) supplemented with 10% Fetal Bovine Serum (FBS) and 1% penicillin-streptomycin. 16HBE14o-cells were seeded at 5.0×10^5^ cells per well of a 6-well plate and incubated at 37°C and 5% CO_2_. After two days, once cells reached approximately 50% confluence, 1 μg of reporter was transfected using Lipofectamine 3000. For co-transfection with U1 expression plasmids, a 1:50 ratio was used. Total RNA was extracted from cells 48 hours post-transfection using TRIzol (Invitrogen). Samples were treated with 5 units of DNase I at 37°C for 30 min and then extracted with Phenol-Chloroform. For the transfection with different U1 concentrations, the reporter was kept at a constant 20 ng as the amount of U1 added was increased.

HEK293 cells (ATCC) were maintained in MEM supplemented with 10% FBS, 1% L-glutamine, and 1% penicillin–streptomycin at 37 °C in a humidified incubator with 5% CO_2_. One day prior to transfection, 1.5 × 10^5^ cells were seeded into each well of a 24-well plate. Transient transfection was performed using Lipofectamine 3000 (Invitrogen, L3000008) according to the manufacturer’s protocol. Three reporter plasmids (wild type, 3120+1G>A, and 3120G>A) and three treatment plasmids (empty pUC13, wild type U1, and ExSpeU1). 500 ng of each fluorescence/luciferase reporter plasmid was co-transfected with 100 ng of each treatment plasmid, resulting in nine distinct co-transfection conditions. All conditions were processed in parallel.

### RT-PCR splicing analysis

A two-step RT-PCR was performed. cDNA was synthesized using 200 ng RNA and 10 U of AMV RT (Invitrogen). Samples were incubated at 25°C for 5 min, followed by 42°C for 60 min, then 85°C for 5 min. For PCR, half of the cDNA and primers specific to exons 1 and 3 were used. For PCR reactions, initial denaturation was 94°C for 4 min, followed by 94°C for 1 min, 57°C for 1 min, 72°C for 1 min for 22 cycles, then final extension at 72°C for 5 min. Products were resolved on 8% acrylamide gels, stained with Gel-Red (ThermoFisher) and imaged using the Bio-Rad Gel Doc EZ Imager.

### Calculation of exon 18 PSI

The RT-PCR products were analyzed using TapeStation 4150 (Agilent Technologies) housed in the Genomics Core at Case Western Reserve University School of Medicine. The concentrations of the RT-PCR products were measured. To determine percent spliced in (PSI), the calculated concentration of the exon inclusion products was divided by the sum of the calibrated concentrations of the inclusion and skipping products. The resulting proportion was multiplied by 100 to get the percent. PSI averages and standard deviations (n=3) were also calculated for each experiment.

### Imaging and bioluminescence quantification

HeLa cells were imaged 48 h post-transfection using an inverted fluorescence microscope (Axio Observer 7, Zeiss). Following fluorescence imaging, Akalumine-HCl substrate was added to each well at a final concentration of 50 uM. Bioluminescence scans were acquired using an In Vivo Imaging System (IVIS) spectrum scanner (PerkinElmer). The bioluminescent images were processed and quantified using LivingImage software (PerkinElmer).

## ACKNOWLEDGMENTS

We thank every member in the Lou laboratory for their intellectual input. We acknowledge the valuable services provided by the Genomics Core of the CWRU School of Medicine. The human U1 snRNA clone was a gift from Dr. Joan Steitz’ lab at Yale University.

We thank Anton Blatnik for his help with RT-PCR analysis. The research was supported by research grants from the Cystic Fibrosis Foundation, LOU24G0 (HL), DRUMM19R0 and DRUMM24R0 (MLD).

## Notes

### Competing Interest Statement

The authors have declared no competing interest.

